# Electron diffraction captures high-resolution structures from *in vivo* protein nanocrystals of *Bacillus thuringiensis*

**DOI:** 10.1101/2025.06.24.661318

**Authors:** Marcus Gallagher-Jones, Robert Bücker, Quentin Delaunay, Nicolas Coquelle, Elena A. Andreeva, Angharad Smith, Ninon Zala, Paul Montmayeul, R. J. Dwayne Miller, Judy S. Kim, Angus I. Kirkland, Jacques-Philippe Colletier

## Abstract

Bacillus thuringiensis is one of the most widely used biopesticides worldwide. This is owing to the highly-specific pesticidal-proteins various strains produce in the form of nanocrystals. Structure determination from such crystals remains difficult because their small size makes them unsuitable for conventional X-ray crystallography. Here we explore two emerging (cryo-) electron diffraction techniques, namely Microcrystal electron diffraction and serial electron diffraction, as tools for studying the structures of these crystals. Using the mosquitocidal protein Cry11Aa as an example, we compare electron diffraction with state of the art results obtained with an X-ray free electron laser. Our work demonstrates that electron diffraction is a viable alternative for structure determination from such challenging crystals, matching previous results obtained with X-ray free electron lasers. We present a workflow based on readily available instrumentation enabling structure determination directly from the crystals grown *in vivo*, unperturbed by dissolution and therefore preserved in their native state.

## Introduction

The bacterium *Bacillus thuringiensis* (*Bt*) produces in the form of naturally-occurring nanocrystals a variety of pesticidal proteins^1,2^. These are used worldwide to control agricultural pests and insect vector populations in an environmentally sound fashion^3,4^. The intracellular growth of the crystals is limited by the boundary of the *Bt* cell membrane, explaining their sub-micron size. The overall mechanism of action of these crystalline proteins lies in their ingestion by insect or nematode larvae^5^, in whose gut the crystals dissolve upon alkaline (insects) or acidic (nematodes) activation. The released proteins are then activated proteolytically enabling their oligomerization and formation of a transmembrane pore^6^, generally upon interaction with a receptor (Cry and Tpp proteins) or the lipidic bilayer (Cyt proteins)^7,8^. Detailed information of the structure-function relationships of these proteins, and notably how the bioactivation takes place, remains incomplete. This is due to the small size of the natural crystals, which makes them unsuited for structure determination by oscillation-based cryo-crystallography methods at synchrotron light sources. Indeed, their limited number of unit-cells translates to both weak diffraction-signal arising from each crystal and high sensitivity to radiation damage. To circumvent this barrier, investigators initially resorted to crystallization of heterologously-expressed protein or dissolution of the natural crystals followed by proteolytic activation and re-crystallisation^9,10^. This strategy has been successful for a limited number of *Bt* proteins^11–13^. However, information was lost regarding the native crystal packing as it could not be assumed that this would be preserved in the (re)-crystallisation process – even more so, when proteins were proteolytically activated before this step. However, how these proteins arrange within crystals is central to understanding their biological activation cascade, as their function critically depends on both efficient crystallization in *Bt* cells and swift dissolution in the target animal’s gut^14^. The ideal approach is thus to solve these protein structures directly from the natural crystals produced by *Bt*. To date, with one exception^15^, this has been attained only using serial femtosecond crystallography (SFX) at X-ray free electron lasers (XFELs), where ultra-short, highly-intense (sub)-µm-focused X-ray pulses are used to decouple the resolution of recordable diffraction from radiation damage^16^. Using SFX, structures could be solved of half a dozen naturally crystalline pesticidal proteins which, despite widespread application, lacked proper structural characterisation. These include *Bacillus thuringiensis* (*Bt*) Cry3Aa^17^, Cyt1Aa^14^, Cry11Aa and Cry11Ba^18^, Cry1Ca18 and Cry8Ba2^19^, as well as *Lysinibacillus shaericus* Tpp49Aa^20^ and the binary complex formed by Tpp1Aa2 and Tpp2Aa2 (formerly known as BinAB)^21^. Many of these studies pursued insights into the protein dynamics that govern crystal dissolution^14,18,21^, with the aim to ultimately engineer crystal properties, and thereby better their pesticidal activity. It was indeed shown that replacement by mutation of just a single atom (over 2700 to 8000) at select interfaces can affect the shape or size of crystals, their dissolution properties, and their pesticidal activities^14,18,21^. These results demonstrate the value of determining *Bt* nanocrystalline protein structures as they naturally occur, with the goal of understanding their mechanisms of action. However, SFX still presents two weaknesses. Firstly, the method suffers from a high rate of sample consumption. Each crystal is destroyed upon a single exposure, and the fraction of crystals that actually intercept the beam is vanishingly small — at most one in 100 billion — owing to the ultrashort pulse duration (5–40 fs) which, combined with the XFEL repetition rate (30–2700 Hz), results in the beam being active for only ∼0.02–0.1 ns per second of beamtime^22^. Furthermore, in all of the studies mentioned above, samples were delivered to the beam using a liquid jet, which demands large quantities of material even when jet diameter is reduced through gas-dynamic virtual nozzle focusing^23^ (GDVN; 6–10 µm jet) or nanoflow electrospinning^24,25^ (∼0.5–1 µm jet). Typically, 1–4 hours of injection of a crystal slurry at 0.1–0.5 × 10⁹ crystals/ml, flowing at 1–15 µl/min, are necessary to collect sufficient data to solve a structure by molecular replacement methods. Secondly, XFEL beamtime is increasingly allocated to projects seeking to resolve ultrashort-timescale structural dynamics, such as those underlying redox reactions, photo-induced isomerization and bond cleavage, or excited-state proton and electron transfers^26–28^. The continuation of structural biology research on *Bt* pesticidal proteins therefore requires the development of alternative methods for structure determination.

Electron diffraction (ED) techniques, such as Microcrystal electron diffraction (MicroED)^29–31^, are promising alternatives for the study of nanocrystals. In these methods, also referred to as 3D electron diffraction (3DED)^32,33^, diffraction data are recorded continuously as the crystal is unidirectionally rotated under the electron beam, in a manner analogous to conventional (rotation-based) X-ray crystallography^34^. Owing to the greater interaction cross-section and more favourable ratio of elastic to inelastic scattering events of electrons compared to X-rays^35^, meaningful data can be captured from substantially smaller crystals, as illustrated by the hundreds of structures now solved of small molecules^36–38^, peptides^39–41^ and proteins^42–45^. In practice, determination of peptide and protein structures requires vitrification of crystalline sample at cryogenic temperatures, to mitigate the progression of radiation damage^46^. This increases crystal lifetime in the electron beam, enabling collection of partial datasets before degradation. These partial datasets are indexed, scaled, and merged into a complete dataset, which can then be processed using conventional (X-ray) crystallography software^47^ to determine the structure. Most recently, scanning electron diffraction methods such as 4DSTEM and SerialED have emerged as alternative approaches to capturing electron diffraction data^48^. In these, a near parallel beam is scanned to serially collect still diffraction patterns from tens of thousands of localised grid positions. The main limitation of these techniques lie in the flatness of the Ewald sphere associated with the typically-used electron wavelengths (∼0.02 Å), leaving 3D information mostly absent from the diffraction pattern. As such the indexing of crystal-hits is reliant on the availability of prior crystal symmetry information. Scanning ED techniques have been applied to lysozyme and granulovirus inclusions^49^ as well as radiation sensitive zeolites^50^ and peptides^51^. For the proteinaceous samples, the recorded resolution surpassed that achieved by SFX performed on the same samples, highlighting the potential of these approaches in protein nanocrystallography.

Here, we have used MicroED and SerialED as tools for determining structures from naturally-occurring crystals of *Bt* pesticidal proteins. The structure of the mosquitocidal protein Cry11Aa was solved using both methods, and resulting models compared to one another, as well as to that previously obtained by SFX^18^. The potential and limitations of ED were further explored by capturing diffraction data from a variety of *Bt*-grown nanocrystals, including some that had proven too challenging for solution by SFX.

## Results

### MicroED of Cry11Aa

The potential of electron diffraction to solve structures from naturally-occurring *Bt* crystals was first explored by applying MicroED to Cry11Aa. Prior AFM studies of isolated Cry11Aa crystals^18^ demonstrated the average dimensions of the crystals were 800 nm in length, 500 nm in width and 175 nm in thickness, placing them in the ideal size range for MicroED^52^. The size of the protein (72kDa) and unit-cell (a=51; b=110; c=150 Å, α=β=γ=90°) are however challenging for the method. Frozen-hydrated CryoEM grids of isolated crystals were prepared by back-blotting using a 1000-fold dilution of the same crystalline slurry used at the XFEL^18^, resuspended in 10% glycerol. We tested several different grid types but found that holey carbon grids give the best distribution of crystals as the variable hole sizes on the grid accommodates the variety of crystal sizes present in solution (Fig. 1a). Sample grids were loaded into an electron microscope operating at 200 kV and 80 K, and the imaging of crystals in TEM mode revealed the presence of a clear lattice suggesting that the crystals could produce high-quality diffraction (Fig. 1b, c). Samples were first screened visually in diffraction mode by overfocussing the diffraction pattern to form an image. Crystals were centred within the selected area aperture, and the beam was refocussed to form a diffraction pattern. Well-diffracting crystals were selected for further rotation data collection.

**Fig. 1.**
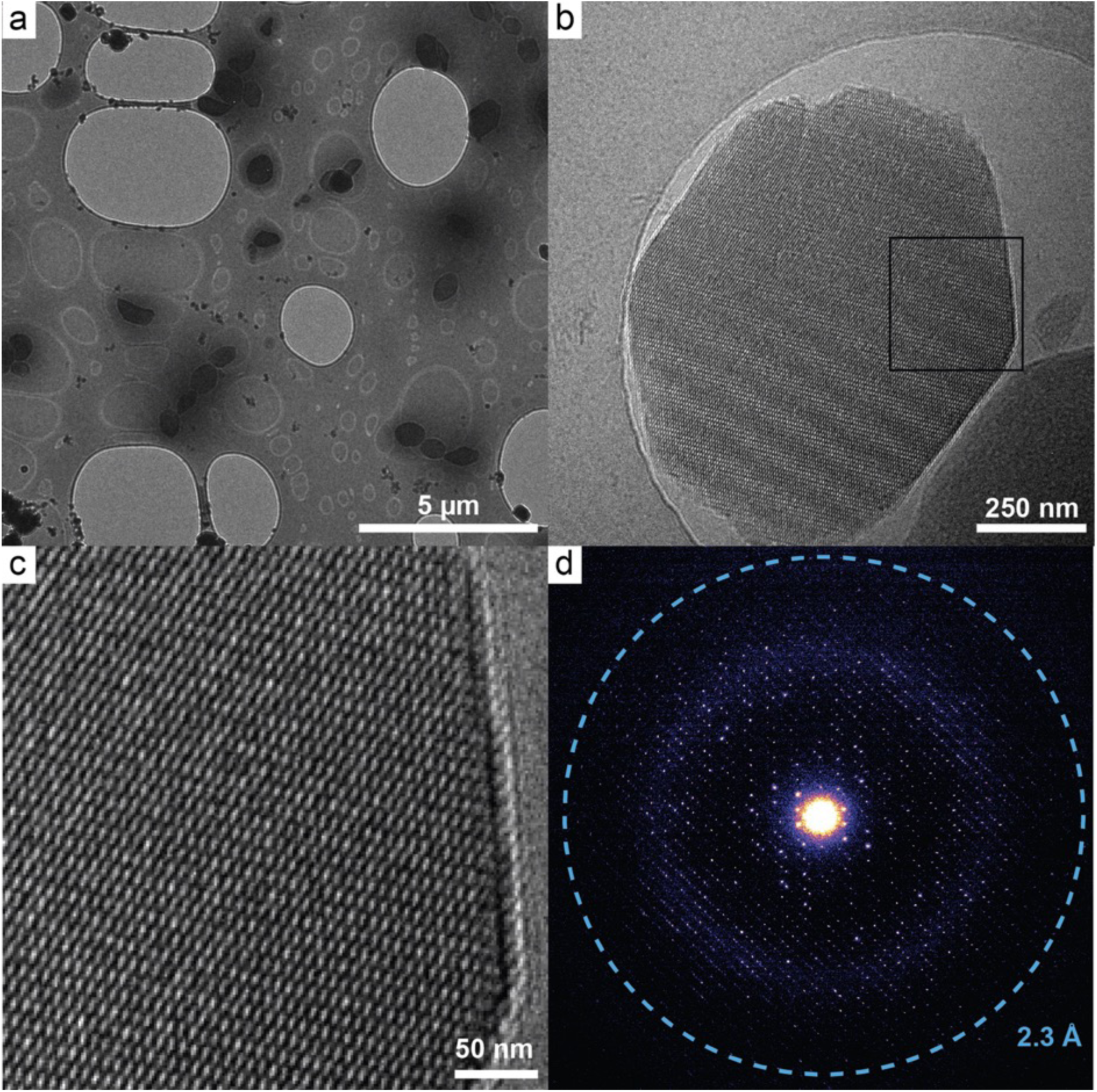
Energy filtered Electron Diffraction of Cry11Aa nanocrystals. a) Overview of frozen-hydrated Cry11Aa nanocrystals on a holey carbon EM grid. b) Higher magnification EFTEM image of an individual crystal showing sharp faceted edges. c) Zoom in of highlighted region in (b) showing clear contrast for individual unit cells. d) Selected area energy filtered ED pattern recorded on a K2 detector in counting mode showing diffraction extending to 2.3 Å resolution.

It has been shown that a monolithic direct electron detector operating in counting mode yields the highest-quality MicroED data^31,53–57^. Thus, given the limited number of unit cells per crystal (<10^5^), we opted to record energy-filtered diffraction data on such a detector (Gatan K2 Summit) to maximize the recoverable signal. Combining a wide-field parallel-beam illumination with a 0.8 µm (in object space) selected-area aperture to define the diffracting region, MicroED data were recorded with signal extending as far as 2.3 Å in favourable cases (Fig. 1d). The sensitivity of crystals to radiation damage was apparent, with diffraction fading over the wedge of data collected, irrespective of the selected crystal. We therefore recorded a small angular wedge of 20 degrees from multiple crystals at a slightly elevated fluence of 3.0 e^−^/Å^2^, corresponding to a cumulated dose of 13 MGy over the whole wedge, well under the experimentally-determined limit of 30 MGy for protein crystals^58^. This was necessary to maximise the signal-to-noise ratio of Bragg peaks at high-resolution, at the cost of some degree of radiation damage. Indeed, while it has previously been shown that it is possible to obtain higher resolution MicroED data by collecting at fluences below 1 e^−^/Å^2 46,54^, this was not possible due to the constraints of our experiment, *i.e.*, a small illumination area and limited number of unit cells, resulting in signal at high-resolution falling below the detection limit of the camera (Supplementary Table 1).

Using this approach, we collected energy-filtered diffraction data from 47 crystals starting from different initial tilt angles. This was necessary to overcome the large variation in diffraction quality of the *in vivo* grown crystals and their preferred orientation on the grids^59^. The quality of data recorded in counting mode was sufficient for unit-cell and space group assignment without the need for additional constraints. For consistency, we indexed all data against the unit cell from the highest-quality dataset, then merged 39 of the 47 datasets into a final set of unique reflections at 2.85 Å resolution (Table 1). The overall completeness of the dataset remained limited to ∼80% owing to the above-mentioned preferred orientation of the plate-like crystals on the EM grid, combined with the limited rotation range of the goniometer. Nevertheless, the SFX derived model provided a successful molecular replacement solution, with an LLG of 58.9 and a TFZ of 4683.6. After several rounds of iterative building and refinement, the final model contained residues 13 to 643 with a R_work_/R_free_ of 0.20/0.24 (Table 1). An overlay with the structure solved by SFX shows a high degree of similarity with an all atom RMSD of 0.79 Å (Fig. 2a). The largest deviations between the structures are seen in the loop (N-ter) and β-turn (C-term) segments adjacent to Cry11Aa β-pin (Fig. 2a). This specific β-strand (residues 341-347), unseen in Cry proteins other than those belonging to the Cry11 primary group, serves as the dimerization interface (C2 symmetry) between Cry11Aa dimers (C2 symmetry), leading to the formation of tetramers (D2) that are the building block (one per unit cell) for Cry11Aa crystal assembly. It is noteworthy that regardless of the technique, the electron density (SFX) and electrostatic potential (MicroED) are well defined for the β-pin itself, while those of the bordering loop (N-ter) and β-turn (C-term) are not (Fig. 2b-d).. This indicates appreciable structural dynamics in these loops at RT (SFX data collected at 293 K) as well as susceptibility to flash-cooling (MicroED and SerialED data collected at 80 K)

**Fig. 2.**
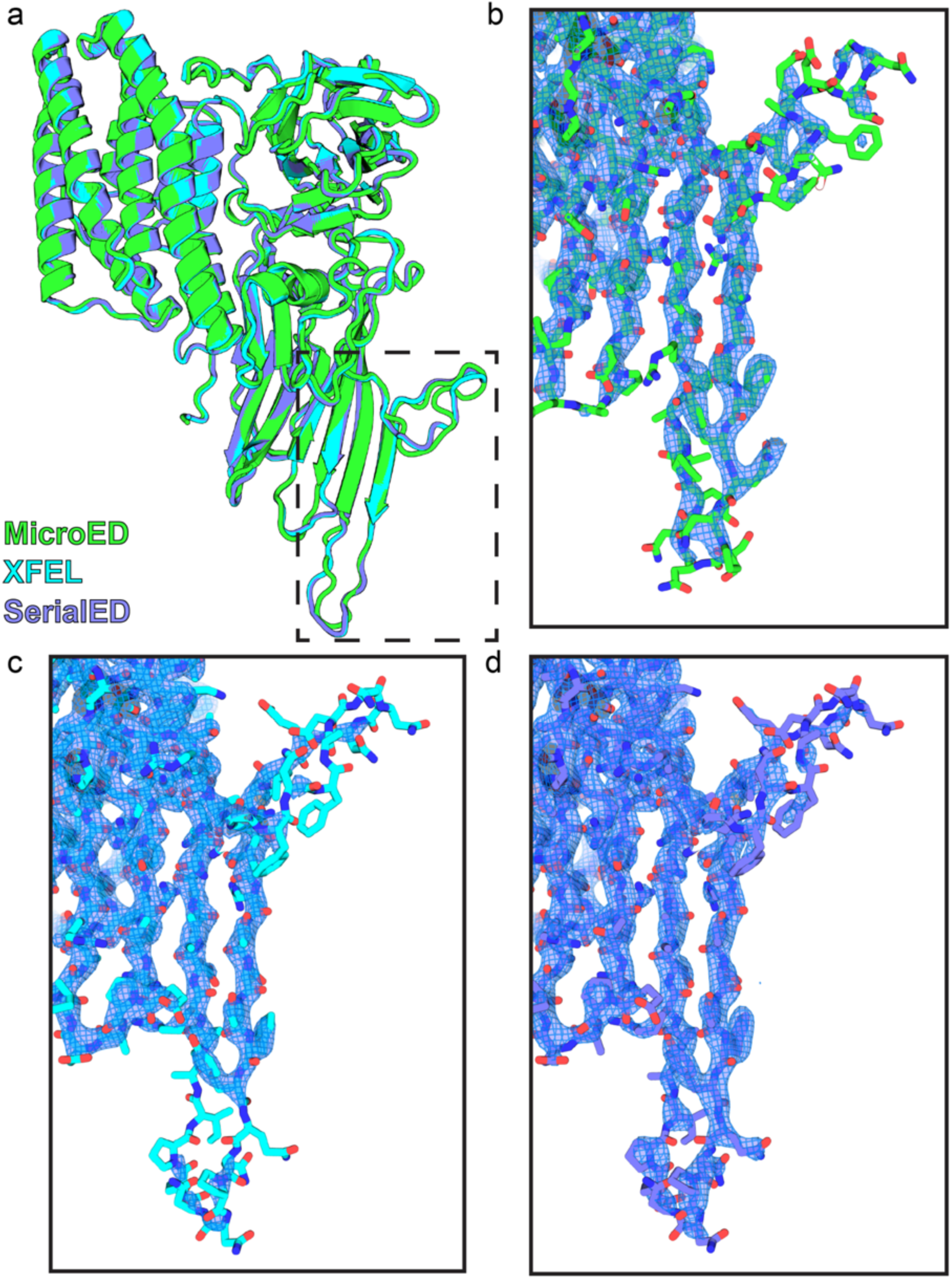
Cry11Aa structures solved by different diffraction methods have the same overall topology and density. a) Alignment of all 3 structures of Cry11Aa in cartoon representation showing good agreement in the overall structure. The dashed box indicate loop and β-turn regions where the majority of the deviations between the structures occur. b) Close up view of the loop and β-turn regions from the MicroED model with 2Fo-Fc coulomb potential map overlaid. c) Close up view of loop and β-turn regions from the SFX model with 2Fo-Fc Electron density map overlaid. d) Close up view of the loop and β-turn regions from the SerialED model with 2Fo-Fc coulomb potential map overlaid. All maps are thresholded at 1.5 σ.

**Table 1.** Data collection and refinement statistics (molecular replacement)

|  | Cry11Aa,<br>MicroED <sup>a</sup><br>energy filtered | Cry11Aa,<br>SerialED <sup>b</sup><br>no 2<sup>nd</sup>-pass scaling | Cry11Aa<br>SFX <sup>c</sup><br>2<sup>nd</sup>-pass scaling | Cry11Aa,<br>SFX <sup>d</sup><br>no 2<sup>nd</sup>-pass scaling |
| --- | --- | --- | --- | --- |
| PDB accession code | 3ZUB | 3ZUK | 7QX4<sup>c</sup> | 3ZUO |
| **Data collection** |  |  |  |  |
| Wavelength (Å) | 0.025 | 0.025 | 1.27 | 1.27 |
| Space group | I222 | I222 | I222 | I222 |
| Cell dimensions |  |  |  |  |
| a, b, c (Å) | 58.1, 153.1,<br>171.7 | 56.9, 155.2,<br>168.0 | 57.6, 155.7,<br>171.1 | 57.6, 155.7,<br>171.1 |
| α, β, γ (°) | 90, 90, 90 | 90, 90, 90 | 90, 90, 90 | 90, 90, 90 |
| Probe size (μm) | 0.8 | 0.1 | 5 | 5 |
| Number of crystals | 39 | 14620 | 48652 | 48652 |
| Number of indexed patterns | 1410 | 12625 | 50613 | 50613 |
| Number of merged images | 1170 <sup>b</sup> | 12625<sup>b</sup> | 48652<sup>b</sup> | 48652<sup>b</sup> |
| Resolution (Å) <sup>a</sup> | 114.25-2.85<br>(2.90-2.85) | 43.5-2.50<br>(2.56-2.50) | 33.55-2.60<br>(2.66-2.60) | 33.55-2.40<br>(2.45-2.40) |
| Number of observations | 739054 (26334) | 4266706 (70735) | 8253629<br>(365007) | 10103779<br>(427713) |
| Number of unique reflections | 15107 (731) | 23881 (1299) | 24198 (1583) | 30621 (1989) |
| I/σ <sup>a</sup> | 10.0 (1.7) | 4.60 (0.99) | 9.50 (1.16) | 7.06 (1.03) |
| Rsplit (%) <sup>a</sup> | n.a. | 0.262 (1.261) | 0.107 (0.954) | 0.119 (1.111) |
| Rpim | 0.084 (0.305) | n.a. | n.a. | n.a. |
| Rmerge | 0.574 (1.830) | n.a. | n.a. | n.a. |
| CC<sub>1/2</sub> <sup>a</sup> | 0.980 (0.214) | 0.932 (0.239) | 0.993 (0.377) | 0.990 (0.304) |
| Completeness (%) <sup>a</sup> | 82.17 (79.80) | 91.16 (77.09) | 99.9 (100.0) | 99.9 (100.0) |
| Multiplicity <sup>a</sup> | 48.9 (36.0) | 178.7 (54.5) | 341.09 (230.5) | 329.96 (215.0) |
| Diffraction limit along a* | 2.6 | 2.5 | 2.5 | 2.3 |
| Diffraction limit along b* | 3.5 | 2.8 | 2.7 | 2.6 |
| Diffraction limit along c* | 2.6 | 2.4 | 2.5 | 2.3 |
| **Refinement** |  |  |  |  |
| Resolution (Å) <sup>a</sup> | 2.85 | 2.50 | 2.6 | 2.4 |
| Number of reflections <sup>a</sup> | 15012 (1293) | 23185 (2109) | 24196 (2247) | 30617 (2590) |
| R<sub>work</sub> / R<sub>free</sub> <sup>a</sup> | 0.20 (0.31) / 0.24<br>(0.40) | 0.28 (0.41) / 0.31<br>(0.48) | 0.17 (0.28) / 0.24<br>(0.37) | 0.20 (0.40) / 0.24<br>(0.44) |
| Number of atoms |  |  |  |  |
| Protein | 5033 | 5085 | 5080 | 5080 |
| Water | 157 | 141 | 261 | 261 |
| B-factors (Å<sup>2</sup>) |  |  |  |  |
| Protein | 38.9 | 39.8 | 50.91 | 50.84 |
| Water | 43.3 | 27.2 | 46.17 | 54.08 |
| R.m.s,d Bond lengths (Å) | 0.003 | 0.003 | 0.004 | 0.002 |
| R.m.s,d Bond angles (°) | 0.560 | 0.490 | 0.633 | 0.425 |
| Ramachandran favored (%) | 96.02 | 96.34 | 95.23 | 96.82 |
| Ramachandran allowed (%) | 3.66 | 3.50 | 4.29 | 2.86 |
| Ramachandran outliers (%) | 0.32 | 0.16 | 0.48 | 0.32 |
| Clashscore | 11.09 | 6.65 | 3.45 | 2.29 |
| Rotamer outliers | 0.37 | 3.61 | 3.68 | 2.00 |

<sup>a</sup> Values in parentheses are for highest-resolution shell. <sup>b</sup> The number of merged images corresponds to number of crystals for serial data and number of frames for MicroED data. <sup>c</sup> Statistics for the SFX data published in previous work (Tetreau et al., 2022, Nat Comm). The corresponding model was used to solve the MicroED and SerialED structure by molecular replacement. <sup>d</sup> Statistics for the SFX data after remerging the original data without the 2<sup>nd</sup>-pass scaling option (`--scale`) in *process-hkl*, and for the SFX model refinement of the deposited structure using 1 cycle of rigid-body and 10 cycles of coordinates and B-factor refinement. All B-factors were reset to 25 Å<sup>2</sup> before refining the structure.

Energy-filtering of the diffraction signal was implemented to reduce the large background caused by inelastic scattering from the bulk solvent and in this manner to improve the signal-to-noise ratio (SNR) of the data^45,60,61^. Whether it was strictly necessary for structure determination, however, remained an open question. To address this, and to assess the benefits of energy-filtering for MicroED data collected on Cry11Aa nanocrystals, we recorded small 5-degree wedges (+/-2.5 degrees) of data with and without the energy filtering, alternating the order of filtered and unfiltered data collection (Fig. 3a,b). Given the isotropic nature of the diffuse background from inelastic scattering, we subtracted a radially-averaged background from the unfiltered data to bring the two datasets onto the same scale for comparison. In line with previous reports^57^, we found that regardless of the order in which data are collected, the SNR of individual Bragg peaks is generally higher in the energy filtered data (Fig. 3c)^57^. This improvement was, however, less pronounced than in previous studies, which we attribute to the thin nature of the Cry11Aa crystals (∼170 nm): thinner crystals produce less inelastic scattering, particularly at low tilts, leaving less background for the filter to remove. To verify this hypothesis while evading the caveat that radial subtraction corrects the background offset but not the underlying counting statistics, we went on collecting 29 additional datasets in the same conditions (20° wedges, 3.0 e-/Å^2^ fluence) yet without energy filtering. The dataset merged from 27 of these extended to 3.73 Å resolution (Table 2), pointing to a dramatic loss in resolution. This loss could be mitigated by excluding datasets where the stage had started from a higher tilt angle— *i.e.*, those where electrons would pass through more material and thus have a higher chance of inelastic scattering. This exclusion strategy resulted in a merged dataset at 2.99 Å but at a severe penalty to completeness (Table 2). These results highlight the necessity of energy filtering for obtaining complete, high-resolution data from Cry11Aa nanocrystals.

**Fig. 3.**
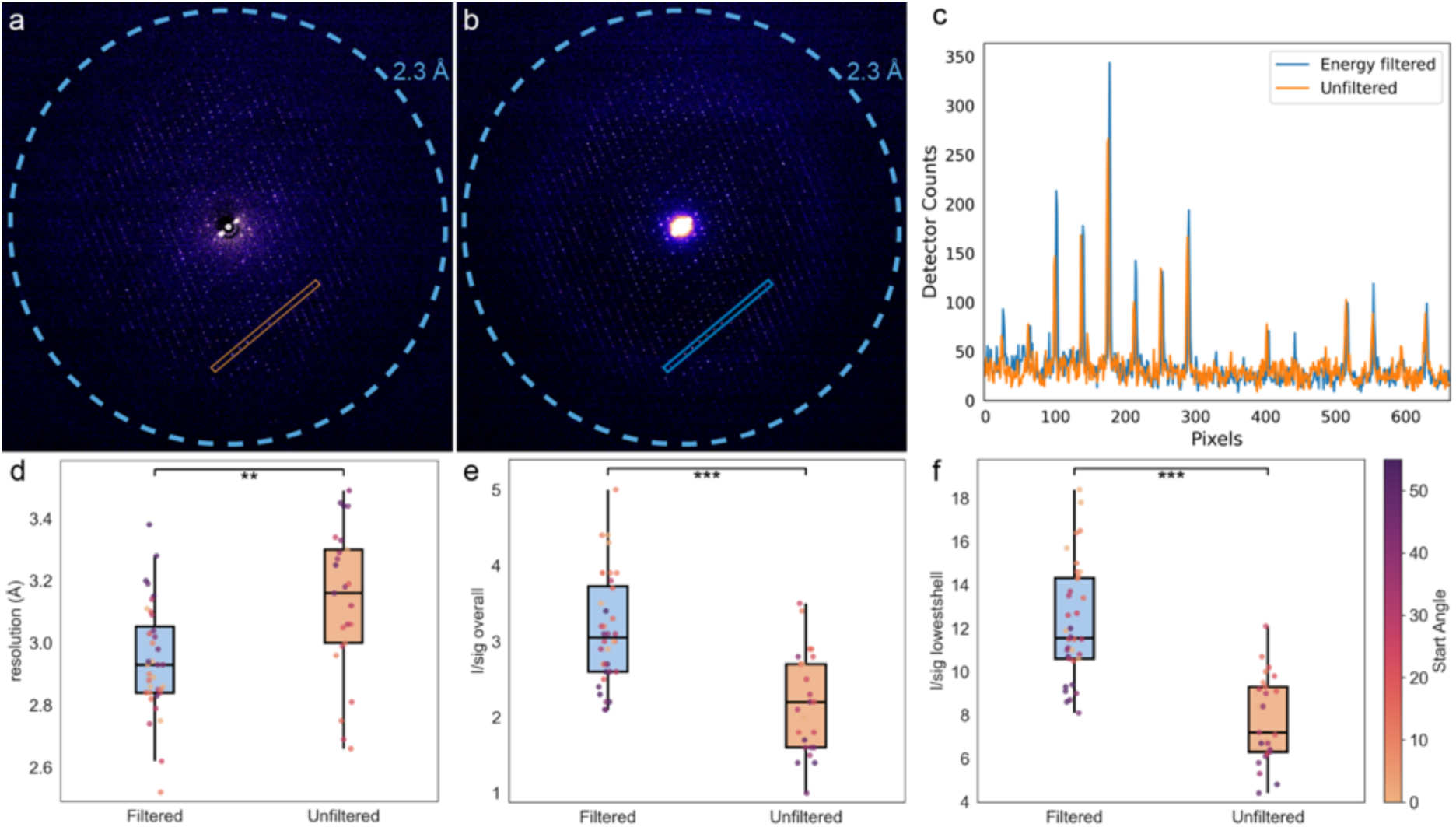
Energy filtered electron-counted MicroED data shows significant improvements in data quality compered to unfiltered data. a) Summation of a five-degree wedge MicroED dataset of Cry11Aa without energy filtering. A radial average background has been subtracted to remove the influence of the inelastic scattering b) Summation of a five-degree wedge of energy filtered data collected from the same Cry11Aa nanocrystal after (a) was collected. A 30 eV width energy slit was used for this dataset. c) Line profile of the peaks indicated in a and b. d - f) Box plot of the final resolution (d) overall I/sigma (e) and I/sigma in the low-resolution shells (f) of individually processed datasets with and without energy filtering. Individual data points are shown and are coloured according to the angle at which rotation was started when data was collected. Stars indicate P values < 0.05 (*), < 0.01 (**) and < 0.001 (***).

**Fig. 4:**
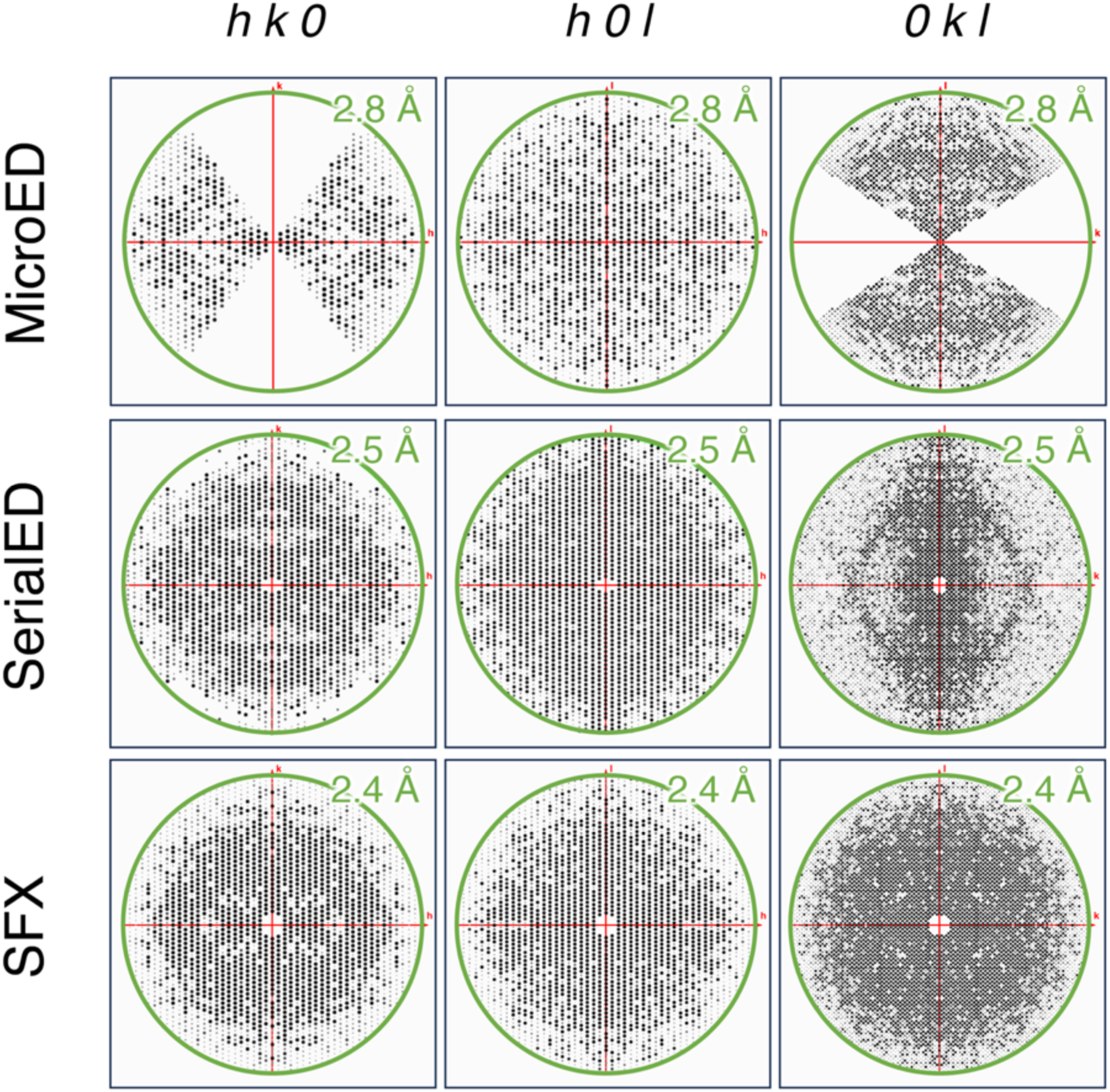
Resolution and completeness of data vary depending on the nanocrystallographic structure determination approach. Integrated reflections are shown for the hk0, h0l and 0kl zones in each of the three datasets. The MicroED data extend to 2.8 Å after merging, A large wedge of reflections is missing due to the limited rotation range of the goniometer installed in the microscope. The SerialED data extend to 1.9 Å, but are only isotropic to 2.5 Å due to preferential orientation of the Cry11Aa crystals on the grid. The SFX data extend to 2.4 Å resolution and are fully isotropic.

**Table 2.** Data collection statistics for MicroED data without Energy-filtering.

|  | Cry11Aa, MicroED<br>- unfiltered | Cry11Aa, MicroED<br>- unfiltered, high-res cutoff |
| --- | --- | --- |
| **Data collection** |  |  |
| Wavelength (Å) | 0.025 | 0.025 |
| Space group | I222 | I222 |
| Cell dimensions |  |  |
| *a*, *b*, *c* (Å) | 58.0, 153.1, 171.6 | 58.0, 155.5, 171.8 |
| $\alpha$, $\beta$, $\gamma$ (°) | 90, 90, 90 | 90, 90, 90 |
| Probe size (μm) | 0.8 | 0.8 |
| Number of crystals | 27 | 27 |
| Number of indexed patterns | 810 | 810 |
| Number of merged images | 450 | 330 |
| Resolution (Å)<sup>a</sup> | 85.81-3.73<br>(3.79-3.73) | 85.91-2.99<br>(3.05-2.99) |
| Number of observations | 123801 (6442) | 141536 (2803) |
| Number of unique reflections | 7255 (364) | 10362 (261) |
| *I*/σ<sup>a</sup> | 6.8 (3.7) | 5.2 (2.0) |
| R<sub>pim</sub> | 0.123 (0.209) | 0.146 (0.320) |
| R<sub>merge</sub> | 0.481 (0.814) | 0.544 (1.013) |
| CC<sub>1/2</sub><sup>a</sup> | 0.743 (0.373) | 0.940 (0.388) |
| Completeness (%)<sup>a</sup> | 87.10 (87.90) | 61.77 (34.51) |
| Multiplicity<sup>a</sup> | 17.1 (17.7) | 13.7 (10.7) |

<sup>a</sup> Values in parentheses are for highest-resolution shell.

As a final assessment of where energy-filtering makes the biggest difference, each of the diffraction datasets was analysed individually, applying the same data quality cutoff of CC½ > 0.3. Data reduction statistics were thereafter compared via Welch’s t-test. We find that the energy filtered data (n = 39) are significantly better than the unfiltered data (n=27) in several key indicators including resolution (*P* = 0.0015), overall I/σ (*P* = 2.2 e^−6^) and I/σ at low resolution (*P* = 5.1 e^−10^) (Fig 3d-f). This difference is more pronounced when the starting angle of the stage is far from zero, *i.e.*, when the apparent thickness of the sample is greatest, supporting the above hypothesis that the thinness of Cry11Aa crystals is the reason for their reduced inelastic scattering at low tilts. This is particularly obvious when looking at I/σ in the low spatial frequencies which suffer greatly from the increased background due to the inelastic scattering of the solvent (Fig. 3f).

As the contribution of inelastic scattering is isotropic and predominantly additive, we finally inquired whether the alternative approach of subtracting the inelastic background would have yielded data of similar quality as those obtained with energy-filtering. We found this approach to be sub-optimal, with the SNR after background subtraction being reduced compared to that of energy-filtered data (Fig. 3c). This is consistent with previous observations that energy-filtering improves the SNR of ED data generally and at all spatial frequencies^60^. Additionally, as energy-filtering reduces the number of electrons reaching the detector, there should be a reduction in coincidence loss at a given flux for electron counting detectors. This will produce more reliable signal in regions where inelastic scattering is high – for example, at lower spatial frequencies, as exemplified by the Cry11Aa MicroED data.

### SerialED of Cry11Aa

The MicroED data were ultimately limited in the highest resolution that could be achieved owing to the need to fractionate a fixed electron fluence over multiple frames. In an attempt to ameliorate this, whilst also reducing cumulative damage, we collected electron diffraction data serially, *i.e.*, with a single high-fluence exposure per crystal, and no rotation of the goniometer during collection^49,62^. For a well-ordered crystal, it should be possible to produce higher-resolution date by expending the entire fluence budget in a single frame. For a diffraction pattern collected serially, 7 – 10x more electrons will reach the detector plane making weaker, high-angle Bragg peaks more visible due to the enhanced per-image SNR. For these SerialED experiments, Cry11Aa crystals were vitrified on CryoEM grids in a similar manner as described above, and loaded into a 200 kV Tecnai F20 (S)TEM equipped with a dedicated diffraction detector^49^. Diffraction data were collected using a ∼110 nm parallel beam raster-scanned across the grids, in a process analogous to serial data collection on solid supports at an X-ray free-electron laser or synchrotron ^63–65^. To maximise data collection efficiency, a low-resolution focussed probe darkfield STEM image was first taken of an area of ∼300 µm^2^ containing tens to hundreds of isolated crystals. Segmentation was performed on this image to automatically map the centroids of each crystal. The microscope was then switched into a parallel nanobeam mode, and a custom scan generator was used to drive the STEM scan coils to position the parallel probe over each crystal. By synchronising this with a hybrid pixel detector with a large dynamic range and fast readout (see Methods), it was possible to record nearly a thousand of such diffraction patterns per hour. In this manner, we collected 14620 SerialED diffraction patterns from Cry11Aa crystals, of which 12625 could be indexed by CrystFEL (v. 0.10.0+9438abb0)^66^ using the *PinkIndexer* algorithm^67^. In practice, all diffraction patterns were collected as 19 doses slices of 0.195 e-/Å^2^ each, but aggregating all available slices proved necessary to obtain the best merging statistics. In what follows, we therefore report on the SerialED dataset merged from data collected at a total dose of 3.7 e⁻/Å² (16 MGy) (Supplementary Fig. 1).

The data extended to a resolution of ∼1.9 Å but were significantly anisotropic (Supplementary Fig. 2). Intensities were initially merged to a more conservative isotropic cut-off of 2.5 Å using a Monte-Carlo approach (Supplementary. Fig. 3), yielding a 92% complete dataset with 78% completeness in the last resolution shell. Given the aforementioned anisotropy we subjected a merged dataset cut to 2 Å resolution to Staraniso^68^ which, using a local <I/sigI> of 1.2 as the diffraction cutoff criterion, found resolution limits of 2.4, 2.8 and 2.4 Å along the a*, b* and c* directions, respectively, and an anisotropy SNR of 6.8. Such an anisotropy of the diffraction was not visible in the SFX data, indicating that it is not an intrinsic property of the crystals but an artefact arising from the preferred orientation of crystals on the grid. In the MicroED data, the issue is largely compensated by rotation of the stage, though a missing wedge in the diffraction plane remains, owing to the limited rotation range of the goniometer and the preferred orientation of the plate like crystals. In the SerialED data, the preferred-orientation problem is instead mitigated by the ∼350-fold larger number of crystals, which increases the chance of sampling a wider range of orientations; this compensation nonetheless remains partial, giving rise to the apparent anisotropy of the merged data which has a slight impact on the electrostatic potential maps in the direction of the missing wedge (Supplementary Fig. 4).

The SerialED dataset was phased by rigid-body refinement using as a starting model the SFX structure. After several rounds of refinement (alternating between reciprocal-space and real-space methods), the final structure had Rwork and Rfree values of 0.27 and 0.31, respectively (Table 1) – notably worse than the MicroED model. Presumably, the issue is not with increased radiation damage, since discarding any of the 19 dose slices collected per crystal resulted in a deterioration of the merged statistics, and Fourier difference maps (see Methods) calculated from ‘early’ and ‘late’ datasets merged from the first five and last five data slices, respectively, showed no signs of specific radiation damage. We thus asked whether it could stem from the comparatively low number of images used to produce the SerialED dataset. Indeed, the published 2.6 Å resolution SFX dataset was originally merged from 50613 diffraction patterns (MC, with second-pass scaling), *i.e.,* a nearly 4-fold increase in multiplicity compared to the SerialED dataset. To test the hypothesis, we merged two new SFX datasets from the publicly available data (https://www.cxidb.org/id-190.html) using the same protocol applied to the serialED dataset (MC, no second-pass scaling), *i.e.*, one comprising all indexed patterns, to a resolution of 2.4 Å; and the other comprising 12,700 randomly-selected indexed patterns, to a resolution of 2.6 Å. Thus, application to the SFX dataset of the same merging protocol used for the SerialED dataset (MC, no second pass scaling) results in better merging statistics than those published initially. Refinement of the published structure against either dataset nevertheless yielded Rfree and Rwork values similar to those published originally (Rwork/Rfree ∼ 0.20/0.24), showing that the origin of the high Rwork/Rfree of the SerialED structure lies not in the number of images used to merge the dataset. Having excluded that the issue originates from completeness (same as the MicroED dataset), multiplicity or radiation damage, we propose that it either stems from heterogeneously vitrified crystals yielding a diversity of subtly different crystalline structures, or from sub-optimal performance of the novel, high dynamic-range Medipix3 electron-detector that was used to collect the SerialED data. Post-refinement of serial data is often key to solving high resolution structures when the number of collected diffraction images is low. It is also useful to discard outlier diffraction patterns whose inclusion in the merged datasets in fact deteriorates the merged statistics. It is thus naturally that we asked whether or not post-refinement of the SerialED data would have yielded a higher quality dataset. All partiality models and merging options available in the *partialator* module of crystFEL were tested. The *ggpm* model was found to be most suitable, which yields iterative improvements across all diffraction data quality metrics (I/σI, Rsplit, CC*, CC1/2) (Table 3). This improvement comes at the cost of data completeness, as not only reflections but full patterns are rejected at each post-refinement iteration. The optimal compromise was achieved with 10 iterations (5.5 % of crystals rejected), producing a dataset with 82% overall completeness at 2.5 Å resolution and 58% completeness in the outermost resolution shell. Refinement against this dataset of the Cry11Aa structure determined on the basis of the MonteCarlo dataset resulted in a 0.5% reduction in both Rwork and Rfree (from 0.21/0.26). This drop was ∼1.5% when refining the Cry11Aa structure against a dataset produced after 50 iterations (21.9 % of crystals rejected) and characterized by 67% overall completeness (and 39 % completeness in the outermost resolution shell), but dramatically improved reflections statistics (see Table 3). These findings suggest that, in some cases, prioritizing higher-quality reflections over completeness can be beneficial. However, since our goal is to compare diffraction data and structures from MicroED and SerialED, we below focus on the SerialED dataset processed with Monte Carlo averaging, which not only aligns better with the merging protocol used for the published SFX dataset but ensures a closer match in completeness to the MicroED dataset.

**Table 3.**
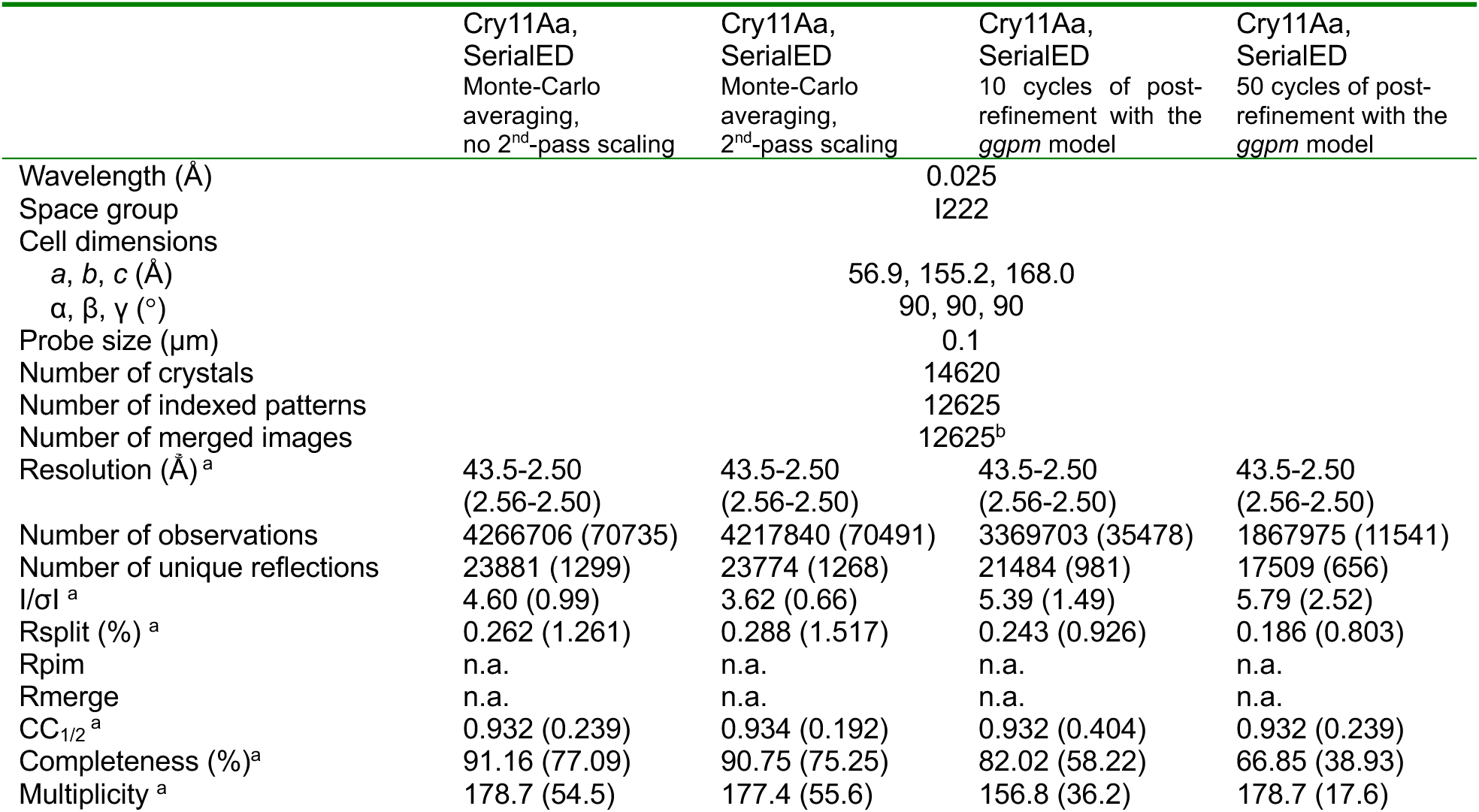
Data collection statistics for Monte-Carlo averaged and post-refined SerialED datasets.

With an aim to provide insight into the effects of cumulative e-damage (MicroED vs. SerialED), vitrification to 80 K (SerialED vs. SFX) and their confluence (MicroED vs. SFX) on the Cry11Aa structure, we interrogated the MicroED and SerialED models using Difference (Cα)-Distance Matrices (DDMs)^69^ (Fig 5a). In the DDM computed between the MicroED and SerialED models, it can be seen clearly that at the monomer level, cumulative electron damage is manifest as an increase in the distance between the fully α-helical domain I and the β-pleated domains II and III. These changes result in a compaction of the two dimers that associate to form the unit cell tetramer, with the dimerizing domains III nearing one another by c.a. 3.5 Å. At variance, the DDMs computed between the SFX model and the MicroED or SerialED models both show a global compaction of the protein, consistent with the ∼3.5% reduction observed in unit cell volume and attributed to the effect of vitrification to 80 K. In both the MicroED or SerialED models, this is seen as domain I nearing domain II which itself nears domain III, consistent with the alignment of the three domains along the b-axis showing the greatest sensitivity to temperature. The compaction of the seven α-helix bundle composing domain I is also visible in both DDMs, to the extent that helices can literally be counted. In the SerialED-SFX DDM, the nearing of domain I and domain III is also visible, which again is expected given the compaction of b-axis. That this nearing is not seen in the MicroED-SFX DDM is likely the result of the change induced by cumulative electron damage.

**Fig. 5:**
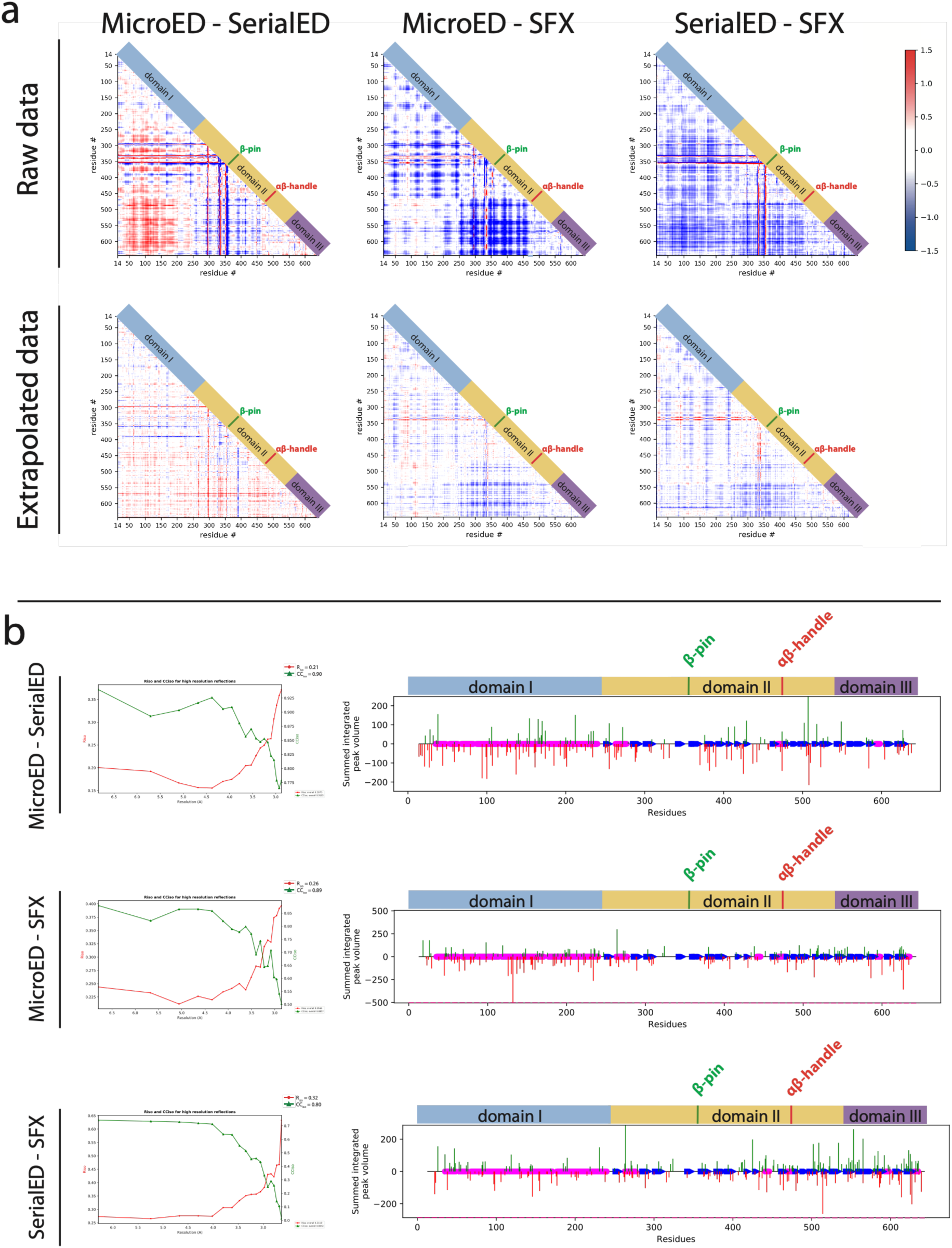
Models derived by MicroED, SerialED and SFX show subtle strucural differences that point to increased disorder in the MicroED and SerialED datasets compared to the SFX dataset. a) Difference (Cα)-distance matrices comparing the MicroED and SerialED Cry11Aa structures (left), the MicroED and SFX structures(middle), and the SerialED and SFX structures (right), respectively. b) The three compared datasets are isomorphous to one another (left panels), making the calculation and interpretation of Fourier difference maps a valuable tool to obtain insights into structural differences. In the three cases (MicroED vs. Serial ED; MicroED vs. SFX; SerialED vs. SFX), we observe larger loss of electron density (negative peaks) than gain of electron density (positive peaks) indicating increasing disorder in the crystals upon cumulative electron damage or vitrification to 100 K. Also in the three cases, we see more peaks on domain I than on the other domains. We note, however, that larger peaks are seen on domain III in the Fobs^SerialED^-Fobs^SFX^ map.

To further investigate the structural differences between the MicroED, SerialED and SFX models, we used Xtrapol8^70^, a software that computes Fourier difference maps between two datasets, providing insights into the structural regions which are prone to structural rearrangements, as well as generates extrapolated structure factors to identify and model low-occupancy intermediate-states which might be present between these two datasets (Fig 5b). All three Fourier difference maps (namely, Fobs^MicroED^-Fobs^SerialED^, Fobs^MicroED^-Fobs^SFX^, Fobs^SerialED^-Fobs^SFX^) contained features above 3.2 σ. Models derived from refinement against extrapolated structure amplitudes show reduced global deviation from their reference state, from which we deduce structure factor extrapolation is a convenient procedure to examine the local effects of vitrification and/or radiation damage decoupled from global changes (inter-domain movements) arising e.g. from unit cell compaction. Taken together, the data illustrate that the SerialED structure shows less hallmarks of cumulative radiation damage inferred from reduced global changes in the protein structure. This, combined with the higher SNR of individual datasets results in data of higher resolution than could be achieved with MicroED. Nevertheless, SerialED shares with MicroED the requirement that crystals are vitrified before data collection, which we show induces global structural changes in the crystalline protein.

### SerialED of other *Bt* crystalline-toxins

The improvement in resolution observed with Cry11Aa encouraged us to test the limits of our approach by collecting SerialED data from several other *Bt* crystalline-toxins. A different microscope setup was used in these endeavours, reflecting changes in available equipment. CryoEM grids were prepared for three types of crystalline-toxins, viz. *Bt israelensis* Cyt1Aa, *Bt alesti* Cry1Ae and C11AB, a fusion of *Bt israelensis* Cry11Aa with the 77-residue C-terminal extension of *Bt jegathesan* Cry11Ba^18^. A model currently exists of Cyt1Aa, obtained from SFX data collected at 1.8 Å resolution^14^, while a model for Cry1Ae has yet to be obtained. The crystals of the C11AB fusion were previously analysed by SFX but only diffracted to 6 Å at EuXFEL^18^ . It remained unclear whether this is due to their small size (< 500 nm), making them too challenging even for SFX, or to intrinsic disorder owing to suboptimal packing of the fusion.

In our studies, we first screened purified crystals by SEM and TEM to assess their suitability for ED (Figure. 6a-c, Supplementary Fig. 5). In all cases a lattice could be observed by TEM, although the Cyt1Aa and Cry1Ae crystals appeared to be quite thick due to their bipyramidal morphology. In contrast the C11AB crystals appeared amorphous in the SEM (Supplementary Fig. 5d) but were shown to be small crystalline fragments when viewed by TEM, each with a single lattice visible (Figure. 6c). SerialED datasets were collected on a JEOL GrandARM2 operating at 300 kV from each of the three crystal preparations using a ∼80 nm parallel beam, which in some cases impinged as few as 750-unit cells. Cry1Ae and Cyt1A produced diffraction patterns extending to 2.3 Å, reaching the resolution-limit of the detector in this experimental setup (Figure. 6d-f), while diffraction to 3.0 Å was observed for C11AB. Thus, whilst Cyt1A crystals did not diffract to a resolution comparable to the one achieved with SFX (1.8 Å), diffraction of C11AB crystals showed significant improvement. Given the greater electron transparency of the C11AB crystals as opposed to the Cyt1A and Cry1Ae crystals, we suspect these engineered crystals are closer to the ideal thickness range for ED^52^ (Figure. 6c). These experiments demonstrate the potential of SerialED to extract data at meaningful resolutions from crystals that cannot be exploited by any other diffraction technique.

**Figure. 6.**
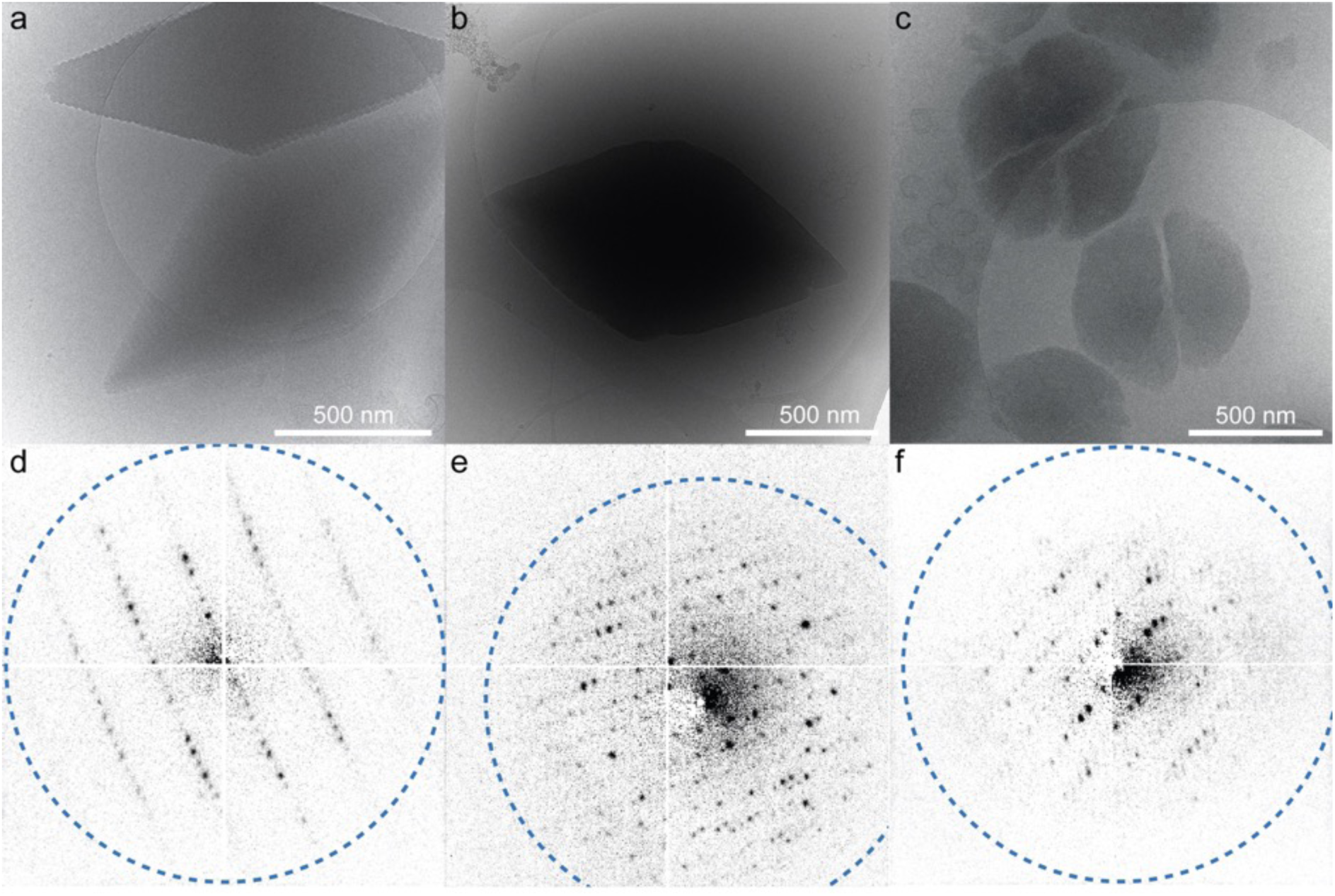
SerialED can capture high-resolution diffraction from a variety of *Bt* derived nanocrystals. a – c) Representative TEM images of crystals of Cry1Ae, Cyt1Aa and C11AB respectively with visible lattice fringes. d - f) Representative SerialED diffraction patterns from crystals of Cry1Ae, Cyt1Aa and C11AB respectively. Blue dashed circle indicates 2.3 Å resolution.

## Discussion

Our results illustrate the potential of electron diffraction methods for solving challenging nanocrystalline protein structures. We here have focused on naturally-occurring nanocrystals of 27-133 kDa pesticidal proteins featuring ∼50,000 – 1,000,000 unit cells. In the natural context, these crystals are produced within the confines of *Bt* cells during their starvation-induced sporulation, explaining that they remain limited in size. Owing to the limited number of unit cells, these crystals pose a challenge for conventional crystallography, which explains why until this work, atomic-resolution structure determination has been limited to scarce beam times at XFEL sources. We show that for crystals of Cry11Aa, featuring 75,000 unit cells on average, electron diffraction data compares favourably with that obtained by SFX, with MicroED and SerialED achieving similar resolution (2.5-2.9 Å). Energy-filtering of the MicroED data was essential to exploit the high-resolution diffraction-signal as the comparatively large amount of solvent surrounding these small crystals produced a significant inelastic background, which furthermore varied along the tilt axis due to changes in the apparent ice-thickness. The improvement in MicroED data quality and resolution observed upon applying energy-filtering is consistent with previous observations on other protein crystals^31,45^. For MicroED, radiation damage remains a persistent problem, requiring the merging of data from many different crystals (here 39) and, alongside the need to fractionate the fluence budget over many frames, limits the achievable resolution. This led us to investigate the potential of SerialED, where a unique high-dose pattern is collected for each of the thousands of irradiated crystals, to maximise the resolution and statistical relevance of the structural data. SerialED allowed diffraction data to be collected at resolutions beyond what was possible by MicroED and even surpassed some of the results obtained by SFX (C11AB). Our results demonstrate that both MicroED and SerialED are well placed to be ideal tools for the study of the structure of *Bt* insecticidal proteins from naturally occurring nanocrystals. Our energy-filtered MicroED data were of sufficient quality to naively assign the correct unit cell parameters from the measured data, making it the ideal choice for studying novel proteins lacking pre-existing crystallographic information. Indeed, MicroED data allowed us to obtain a reasonably complete structure with sufficient resolution to unambiguously assign residues in all key domains of the protein, including the highly mobile loops bordering the β-pin. Unit cell assignment remains challenging for SerialED^71^. Here, the space group assignment remained ambiguous between I222 (correct) and the three alternative lower-symmetry C2 space-groups. With the confident assignment of lattice symmetry based on MicroED, it was possible to collect SerialED data of the same crystals, which merged at a higher resolution by mitigating the impact of cumulative electron damage to the biological structure, and achieved a higher completeness by sampling a larger population of crystals on the EM grid.

Our preliminary analysis of 3 other naturally-crystalline proteins from *Bt*, including a synthetic fusion of Cry11Aa with the C-terminal tail of Cry11Ba (C11AB) that had diffracted poorly at the XFEL^18^, further highlights the potential of SerialED to overcome crystal size and quality limits and be applicable to *Bt* derived as well as morphologically similar crystals. Indeed, we have studied two crystalline proteins where ED achieved a superior resolution compared to SFX. Nevertheless, the highest resolution for Cyt1Aa was still achieved by SFX at XFEL, consistent with results from other studies^59,72^. There can be several reasons for this including the higher intrinsic sensitivity of some proteins to radiation damage by the electron beam^46^ and the morphology of the crystals. In particular, the bipyramidal shape of Cyt1A (and Cry1Ae) was not optimal for ED as the centre of the crystal was too thick for electrons to penetrate. This is particularly problematic for crystals with large unit cell dimensions as it has been demonstrated that a minimum thickness of 4 - 5-unit cells is required to acquire sufficient data for structure determination by MicroED^31,73^. Indeed, among the structures that have currently been solved by MicroED in the PDB we note that there seems to be a link between unit cell volume and the resolution achieved, likely due to the finite thickness accessible to electrons (Supplementary. Fig. 6). Our current work, as well as other recent SerialED work^74,75^, show the potential of this technique to overcome such limits without the need of higher electron accelerating voltages (Supplementary Fig. 6) Irrespectively, our data show that atomic-resolution structural-insights into *Bt* nanocrystalline-toxins are within reach of ED techniques. Of note, it has been reported that it is possible to obtain higher resolution MicroED data by collecting at fluences below 1 e^−^/Å^2 46,54^, but this was clearly not the case here. We found that for crystals featuring a reduced number of unit cells and probed by small electron beams, such as those of Cry11Aa in this study, large doses must be used to collect meaningful data.

Of the hundreds of *Bt* crystalline-toxins, only a handful have been structurally characterised experimentally. Resistance to *Bt* proteins is increasing and being able to experimentally characterise new, functional mutants is critically important. The workflow presented here positions ED as a vital tool in improving our understanding of the native packing of these pesticidal proteins within the bacterial cell. From a broader perspective, ED techniques could also be relevant for similarly sized naturally occurring crystals produced by other biological systems. Indeed, there is a growing body of work demonstrating the advantages of electrons for studying molecular nanocrystals grown within a cellular context^76–78^. The previously mentioned limitation on sample thickness means electron diffraction cannot be considered a general technique for structure determination of all protein nanocrystals grown *in vivo*. Indeed, a current requirement to applying ED techniques is that crystals remain stable in water or another vitrification compatible buffer after extraction from cells, as probing their structure *in situ* would be challenged by the augmented thickness of scattering material. In this context, related electron-microscopy methods such as cryoFIB-SEM are required to enable exposure of the crystals (or fragments thereof) from the cells^79,80^. The work of Yang et al is particularly relevant as the system studied in this paper, eosinophilic major basic protein (eMBP), was approached by SFX, but structural insights were limited to demonstrating that the packing of individual protein chains differs in the natural nanocrystals and those obtained by recrystallising the protein i*n vitro*^81^. By contrast ED techniques combined with cryoFIB-SEM, were able to interrogate the *in-situ* structure of eMBP and assess structural differences arising from eosinophil activation^77^.

In conclusion, our work demonstrates that ED is a valuable addition to the *Bt* structural biology toolbox. Our results suggest that recent (and future) advances in detector technology^55,82,83^ and sample preparation^84–86^ will broaden the applicability of these techniques. More specifically, MicroED provides a robust and accessible platform for protein nanocrystallography using conventional electron microscopes and established crystallographic software. SerialED, by contrast, requires more specialised instruments but has the potential to produce data of higher resolution and/or completeness. Furthermore, we expect incorporating energy-filtering into the SerialED workflow will provide similar benefits to those observed in MicroED, such that data quality may be further improved. As new tools and protocols emerge for reliably producing protein nanocrystals we expect a growth in the application of ED for protein structure determination, similar to the boom that has been observed for chemical crystallography^87^.

## Materials and Methods

### Crystal expression and purification

Cry11Aa crystals were produced and purified as previously described (Tetreau et al, 2022, Nat Comm). Briefly, acrystalliferous (plasmid-curated) cells of *Bacillus thuringiensis israelensis* (strain 4Q7; The Bacillus Genetic Stock Center (BGSC), Columbus, USA) were transformed by electroporation with the pWF53 plasmid^88^ featuring : (i) the *cry11aa* gene under control of its natural promoter; (ii) the *P20* gene under control of the Cry1Ac promoter; (iii) replication origins suited for amplification in *E. coli* and expression in *Bt*, respectively; and (iv) resistance cassettes for erythromycin (*Bt* selection) and ampicillin (*E. coli* selection). Transformed cells were deposited on LB agar medium supplemented with erythromycin (25 µg/mL), and after two days, a colony was swept to inoculate an LB pre-culture, also supplemented with 25 μg/mL erythromycin. For the production phase, precultures were spread on T3 sporulation medium (Tryptose 2 g L-1, Tryptone 3 g L-1, yeast extract 1.5 g L-1, Na2HPO4 1.4 g.L-1, NaH2PO4 1.2 g.L-1, MnSO4, MgSO4, and ionic solution 1 mL per 1 L (in 1 L of H2O: 6.82 g ZnCl2, 101.66 g MgCl2.6H2O, 1.98 g MnCl2.4H2O, 29.4 g CaCl2.2H2O, 13.52 g FeCl3.6H2O, 967 µL 10 M HCl)) containing 25 μg/mL erythromycin., and after incubation for 4 days at 30°C, spore and crystal were recovered using cell scrapers, pooled and resuspended in ultra-pure water. Cells were lysed by several cycles of sonication (1 sec excitation pulse at 0.5 Hz during 1 minute) at 100 W, centrifuged at 4,000 *g* for 45 minutes to remove cell debris and medium, and then centrifuged again at 10,000 g to pellet crystals and spores. The pellet was resuspended in water and the crystals were isolated from the spores and remaining cell debris by ultracentrifugation (23,000 × g, 4°C, 17 h) on a discontinuous sucrose gradient (67-72-79%). After ultracentrifugation, the crystals were recovered at the 67-72% interface, and thereafter washed by several cycles of centrifugation/resuspension in ultra-pure water to eliminate sucrose. The purity and integrity of the crystals was assessed by SDS-PAGE on 12% gels and by scanning electron microscopy (Ultra-55 Zeiss, Zeiss, Germany) on the nano-characterization platform (IRIG-CEA, Grenoble, France), respectively. Crystals were stored in ultra-pure water and at 4°C before use.

Cyt1Aa and C11AB crystals were produced and purified as previously described^14,18^. Transformation, expression and crystal-purification were performed as described above for Cry11Aa.

### Crystal visualisation using scanning electron microscopy (SEM)

Purified crystals of Cry11Aa, Cyt1Aa, Cry1Ae and C11AB were imaged using a Zeiss Ultra 55 scanning electron microscope at the nanocharacterization platform in the Institut de Recherche Interdisciplinaire de Grenoble (IRIG). Samples were coated with a 2 nm thick layer of gold using a Compact Coating Unit-010 sputtering unit (Safematic®) prior to imaging. Images were recorded at 5-20 kV acceleration voltage by collecting secondary electrons using an Everhart-Thornley-Detector (ETD detector) in high-vacuum mode.

### CryoEM grid preparation

For MicroED experiments a slurry of purified Cry11Aa crystals was first diluted 1000-fold in 10% glycerol and 2 µl of diluted crystal suspension were applied to the carbon side of a freshly glow discharged (60 s 20 mA, Pelco Easyglow) holey carbon grid (300 mesh Cu (emResolutions)). A solution of 2 µl of 10% glycerol was then applied to the opposite side of the grid and excess solution was removed by single side back blotting in a Leica EM GP2 (7s, 85% humidity, 4 C) before plunge freezing into liquid ethane. This concentration of glycerol was selected after trialling several different concentrations (5 – 30%) and was found to give the best balance between consistent vitrification whilst preserving a low solvent background.

For SerialED experiments of Cry11Aa, a slurry of crystals was prepared as for the experiment. A volume of 3.5 μl of suspension was applied to the carbon side of a freshly glow discharged hole-array grid (Quanitfoil R1.2/1.3). Two-sided blotting was applied in a FEI Mark IV (FEI Company), before plunge freezing into liquid ethane.

For SerialED experiments of the other *Bt* crystals, crystalline slurries were resuspended in 10% glycerol and 3 µl of these suspensions were applied directly to freshly glow discharged (60s 20 mA, Quorum GloQube Plus) Quantifoil grids (1.2/1.3 400 mesh Cu). Excess solution was removed by double sided blotting in a Vitrobot Mark IV (Thermo Fisher Scientific) using parameters optimised for each crystal type (generally blot time of 10 - 15s, blot force of 10 - 15, 20s wait time, 1s drain time, 80% humidity, 20 c) before plunge freezing into liquid ethane.

### MicroED data collection

Frozen hydrated EM grids were transferred into a JEOL cryoARM300 operating at 200 kV for data collection. The A2 gun lens voltage was reduced from 3.33 kv to 3.13 kv to reduce the incident fluence at the sample. A camera length was selected such that the top edge of the K2 detector corresponded to 2 Å resolution using a gold cross grating as a calibration standard. Diffraction quality crystals were identified in an over-focussed diffraction mode by maximising the excitation of the third condenser lens. Crystals were manually brought to the goniometer eucentric height and the excitation of third condenser lens was then reduced to form a parallel beam with a flux of 0.015 e^−^/Å^2^/s^−1^ for MicroED data collection. Diffraction movies were collected semi-automatically using SerialEM^89^ to synchronise stage movement with the camera operation. Crystals were isolated using a 50 µm selected area aperture (0.8 µm field of view at the sample plane) and each crystal was rotated over a 30-degree wedge with a rotation speed of 0.1 degrees per second giving a total fluence of 4.5 e^−^/Å^2^. Data was recorded on a K2 DE detector in counting mode below an omega filter with a frame time of 1s. Energy filtered data was collected with a slit width of 30 ev or the slits were completely removed for unfiltered data. Due to the low fluence it was possible to record data without a beam stop. A total of 47 zero-loss filtered, and 27 unfiltered datasets were collected.

### Data reduction, phasing and model building for MicroED

Raw data frames in LZW compressed tiff format were pre-processed using a custom python script that performed drift correction, diffraction pattern centring, hot pixel removal, gain correction and binning before outputting data in MRC format. Frames were binned by 2 in the x and y dimensions and 10 in the temporal dimension to give 30 frame stacks where each frame represents a 1-degree wedge of data. Data reduction (spot, finding, indexing, refinement, integration and scaling) was performed in DIALS^90,91^ using the first 20 frames to reduce the impact of radiation damage. A single dataset with the highest quality diffraction was analysed first and the space group was determined to be I222 by the routine dials.refine_bravais_setting. This space group was then used as a constraint during automated processing of all 47 datasets. The xia2.multiplex program^92^ was used to assess the best movies to merge and the final resolution cut off of 2.85 Å using a cluster containing 39 of the 47 datasets. Phasing was performed by molecular replacement using the previously resolved XFEL structure. A single solution was found in Phaser^93^ with space group I222, an LLG of 4683.6 and a TFZ of 58.9. Refinement of this model was performed by alternating between automated refinement in Phenix, using electron scattering factors, and manual adjustments performed in Coot^94^ to produce a final model with an Rwork/Rfree of 0.1950/0.2417.

### Filtered vs. Unfiltered diffraction data background comparison

Pairs of datasets were taken from multiple crystals over a 5-degree angular wedge with the energy defining slits inserted or completely removed. Whether filtered or unfiltered data was collected first was alternated to reduce potential bias due to radiation damage. The raw data was pre-processed using a custom python script that first performed gain correction, binning and summation. In addition, it was noted that the images contained an additional pixel offset between each line in the fast scan direction, likely resulting from additional dark noise due to the sparse nature of the diffraction data (Supplementary Fig. 7). To account for this the median value for each line was calculated using regions of the image that had not encountered any electrons. These median values were used to create a map of pixel offsets per line with the same dimensions as the initial image and this was subtracted from the final data. To compare the unfiltered data on a similar scale to the unfiltered data a radial background was calculated by first identifying the centre of the diffraction pattern and calculating the radial intensity at each resolution bin. This 2D map of radial intensity was then smoothed by a median filter with a kernal size of 20 pixels to reduce the influence of Bragg peaks and finally subtracted from the unfiltered image. Line profiles were drawn across rows of Bragg peaks and the plotted to compare relative intensities for pairs of images.

### SerialED data collection for Cry11Aa

SerialED data from Cry11Aa were collected using a Philips Tecnai F20 field-emission S/TEM operating at 200 kV, equipped with a custom-built hybrid pixel detector (DESY detector group) based on a Medipix3 chip (1556×516 pixels), synchronized to a custom-built beam scanning system (based on a National Instruments PCI-6251 DAQ board) allowing for arbitrary patterns of fast beam movement and scanning. The sample grids prepared as described above were transferred into the S/TEM using a Gatan Model 626 cryo-transfer holder. Detailed Instrument setup, sample screening and data collection are described in detail in^49^. Mapping images over a field of view of 18×18 μm² were collected from each region ADF-STEM mode at a total fluence of <0.1 e^−^/Å². Using image segmentation, positions of nanocrystals are automatically identified for SerialED data collection. For the ensuing data collection step, the beam was defocused using the C2 condenser to give a parallel condition (∼0.1 mrad) with a diameter of ∼110 nm, defined by a 5 µm condenser aperture. By setting the gun lens of the field-emission gun to its lowest excitation and the C1 condenser lens to Spotsize 6, a beam current of 14 pA was achieved with the 5 μm condenser aperture, yielding a flux density of 92 e^−^/A²/s^−1^ on the crystals. Data was collected in a dose-fractionated mode by setting the frame time of the detector to 2 ms with 20 frames collected from each crystal, corresponding to a total fluence (dose) of 0.18 e^−^/A² per movie frame and 3.7 e^−^/A² for the entire movie. During processing, it was found that optimal data quality (gauged by merging metrics and spot intensities) was achieved by summing up all frames for each crystal in the final data processing, i.e., the full fluence of 3.7 e^−^/A² per crystal was used in the final dataset. The projection system of the S/TEM was set to an effective camera length of 3062 mm on the detector, corresponding to a maximum diffraction resolution of 1.8 Å near the long edge of the detector. The camera length and a residual elliptical distortion of ∼2% was calibrated using diffraction ring patterns from a polycrystalline thallous chloride standard (Ted Pella).

### Data reduction, phasing and model building for SerialED

Data were processed using diffracTEM^71^ and CrystFEL (v. 0.10.0+9438abb0)^66^, with PinkIndexer serving as the indexing algorithm^67^. Of 14620 collected images, 12625 could be indexed (*indexamajig* program) and were retained for Monte-Carlo merging (*process_hkl* program). Post-refinement of the data (*partialator* program) was attempted with all models^95,96^, and delivered best results when the *ggpm* model was used (Supplementary Fig. 3). However, completeness and multiplicity were negatively impacted, which resulted in electron potential maps of lesser quality. Likewise, Monte Carlo averaging using the ‘second-pass’ (scaling) option in CrystFEL *process_hkl* yielded data of worse quality, as did the inclusion of polarization corrections or the treatment of Friedel pairs as separate reflections. Hence, reported results are derived from the structure factors amplitudes estimates obtained by simple MonteCarlo averaging, with Friedel law considered true. These issues are likely due to our limited amount of data, which translates to low completeness and multiplicity and therefore to high uncertainty in the estimation of structure factor amplitude. Furthermore, we conjecture that due to the homogeneous size and shape of the crystals, a simple averaging of intensities collected from each one is justified even for a comparatively small number of crystals, as opposed to merging data from more diverse crystals where more involved refinements are beneficial.

Phasing of the SerialED data was achieved by rigid-body refinement using *phenix.refine* within the Phenix crystallographic software suite. Thereafter, iterative cycles were carried out of real-space refinement in Coot^94^ and reciprocal space refinement in *phenix.refine*, until figure of merits plateaued. Analysis of structural differences was carried out using Xtrapol8^70^ and custom-written scripts. Figures were produced with PyMOL and ChimeraX^97^.

In an attempt to further investigate the structural differences between the MicroED, SerialED and SFX models, we used Xtrapol8^70^, a software that computes Fourier difference maps between two datasets, as well as generates extrapolated structure factors to identify and model low-occupancy intermediate-states which might be present between these two datasets (Fig. 5b). For the three pair of datasets, we used k-weighted extrapolated structure factors produced with a k-weight scale of 0.05. Fourier difference peaks shown in Fig. 5b were selected abive a 3.2 σ threshold, integrated to a floor-value of 2.7 σ, and attributed to the closed residue within a 2 Å radius. Negative extrapolated intensities were rescued using the *truncate-and-fill* method in Xtrapol8. All structure underwent 10 cycles of coordinate and B-factor refinement against extrapolated structure factor amplitudes. The extrapolation factors were found to be 3, 2 and 2 for the MicroED vs. SerialED, MicroED vs SFX, and SerialED vs. SFX comparisons, respectively.

### SerialED data collection for other *Bt* crystals

Samples were loaded into a double corrected JEOL GrandARM2 TEM operating in nanobeam diffraction (NBD) mode using a cryo transfer holder (Fischione, Model 2550). Coma and twofold astigmatism were adjusted manually using the ronchigram. With this optical geometry it was possible to form a ∼ 80 nm diameter parallel probe using a 10 µm condenser aperture by adjusting the excitation of the second condenser lens to give Kohler illumination. A simple script was implemented in pyJem to rapidly switch between the convergent STEM illumination and the parallel SerialED illumination. The measured probe current was 22 pA giving a fluence of 0.35 e-/Å^2 per frame assuming a 2 ms dwell time. The camera length was set at 800 mm which corresponds to a true camera length of 1346 mm based on diffraction from the gold cross-grating. Low magnification STEM (2500x magnification with a 512 by 512 raster) was used to identify grid squares with suitable sample distribution, and the stage positions were recorded.

Data collection was performed using a set of scripts implemented in python and Julia to control the movement of the STEM scan coils (Quantum Detectors Scan Engine) and synchronise this with data acquisition on a MerlinEM 4R detector (Quantum Detectors). Data collection proceeded as follows. Firstly, the microscope was set to STEM mode and search STEM scans were recorded at 20kx with an 80 nm step size (i.e. a 128 X 128 scan) and a 20 µs dwell time, giving a fluence of 0.0025 e^−^/Å^2^ per scan. Crystal coordinates were then computationally segmented from the STEM image and the microscope switched to a SerialED mode. The beam was then moved to the segmented coordinates as an arbitrary scan to collect data. Data were recorded as 2ms × 10 frame stacks such that the total fluence at each coordinate is 3.5 e-/Å^2. ‘Good’ hits for further processing were determined based on the amount of signal beyond a low spatial frequency cut-off, to avoid the high signal at the primary beam. A radial background was subtracted to aid with Bragg peak visualisation.

## Data availability

Coordinates and structure factors for Cry11Aa determined by MicroED and SerialED at 2.2Å and 1.9Å resolution cutoffs have been deposited in the RCSB Protein Data Bank under accession codes 9RML, 9RMP and 9RMN respectively. MicroED datasets and SerialED data streams, alongside the microscope control and data analysis scripts, have been placed in a Zenodo repository, entry 15691475.

## Acknowledgments

We thank Brian Federici for kindly providing the pWF53 & pWF45 plasmid used for production of Cry11Aa and Cyt1Aa crystals, respectively. We thank Martin Weik, Didier Lereclus and Michel Gohar for continuous support. We also warmly thank Elke De Zitter for implementation of the use of electron scattering factors in Xtrapol8. MGJ thanks David Waterman for assistance with processing ED data in DIALS. IBS acknowledges integration into the Interdisciplinary Research Institute of Grenoble (IRIG, CEA). This work was supported by the Agence Nationale de la Recherche (grants ANR-17-CE11-0018-01, ANR-20-CE11-0019-02 and ANR-22-CE11-0016 to J.-P.C.). Platform access was supported by FRISBI (ANR-10-INBS-05-02) and GRAL, a project of the University Grenoble Alpes graduate school (Ecoles Universitaires de Recherche) CBH-EUR-GS (ANR-17-EURE-0003). RJDM’s work is Supported by the Natural Sciences and Engineering Research Council of Canada. MGJ, JSK and AIK acknowledge funding from the Engineering and Physical Sciences Research Council to the Rosalind Franklin Institute. QD was funded by the 80PRIME PhD Program of the CNRS and received support from MITACS and the UGA through a Globalink Research Award.

## Notes

RJDM and RB are listed as the inventors of a patent on Serial NED.

