## Supplementary_information for "Electron diffraction captures high-resolution structures from *in vivo* protein nanocrystals of *Bacillus thuringiensis*"

### Supplementary figures

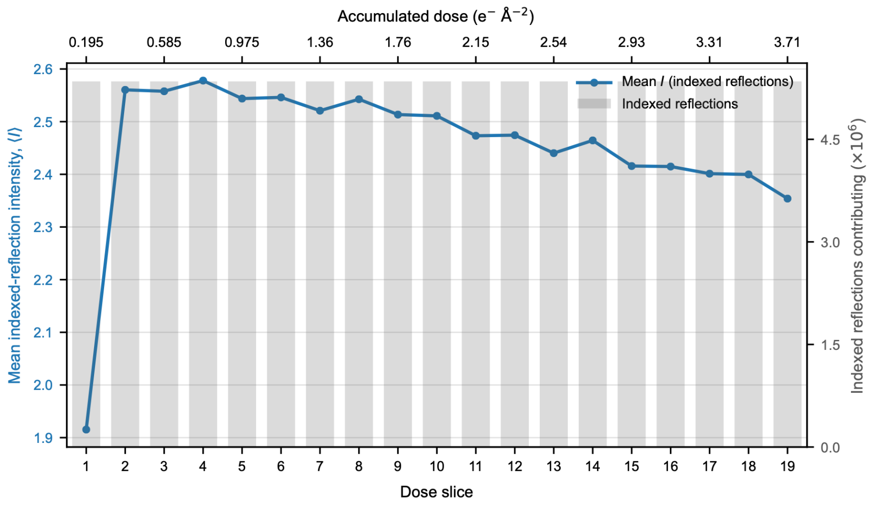


**Supplementary Fig. 1: Dose-dependent decay of Bragg intensities in the Cry11Aa SerialED data.** Mean intensity of all indexed reflections is plotted as a function of dose slice (blue line and circles); grey bars indicate the number of indexed reflections contributing to each mean (right axis). Each dose slice (1-19) corresponds to an electron exposure of 0.195 $e^{-}$ $Å^{-2}$ (0.894 MGy), giving a total accumulated dose of 3.705 $e^{-}$ $Å^{-2}$ (16 MGy). The progressive decrease in mean reflection intensity provides a measure of electron-beam-induced radiation damage during data acquisition.

**Commented [JPC1]:**

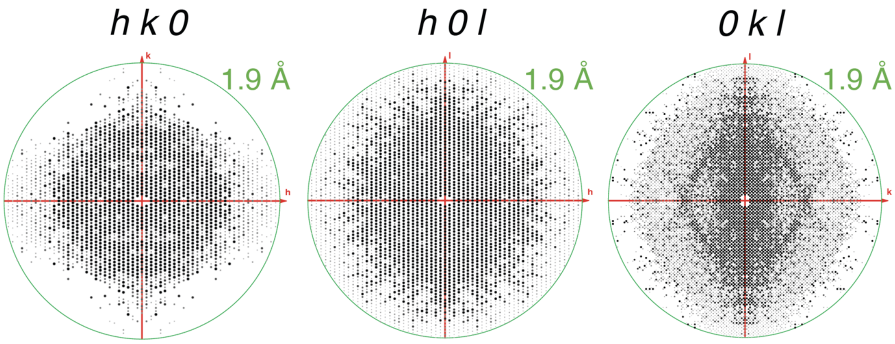


**Supplementary Fig. 2: SerialED data extends to high resolution but shows severe anisotropy. I**ntegrated reflections are shown for the hk0, h0l and 0kl zones for the full range of resolution of the SerialED data**.** Signficant anisotropy is evident in hk0 and 0kl zones.

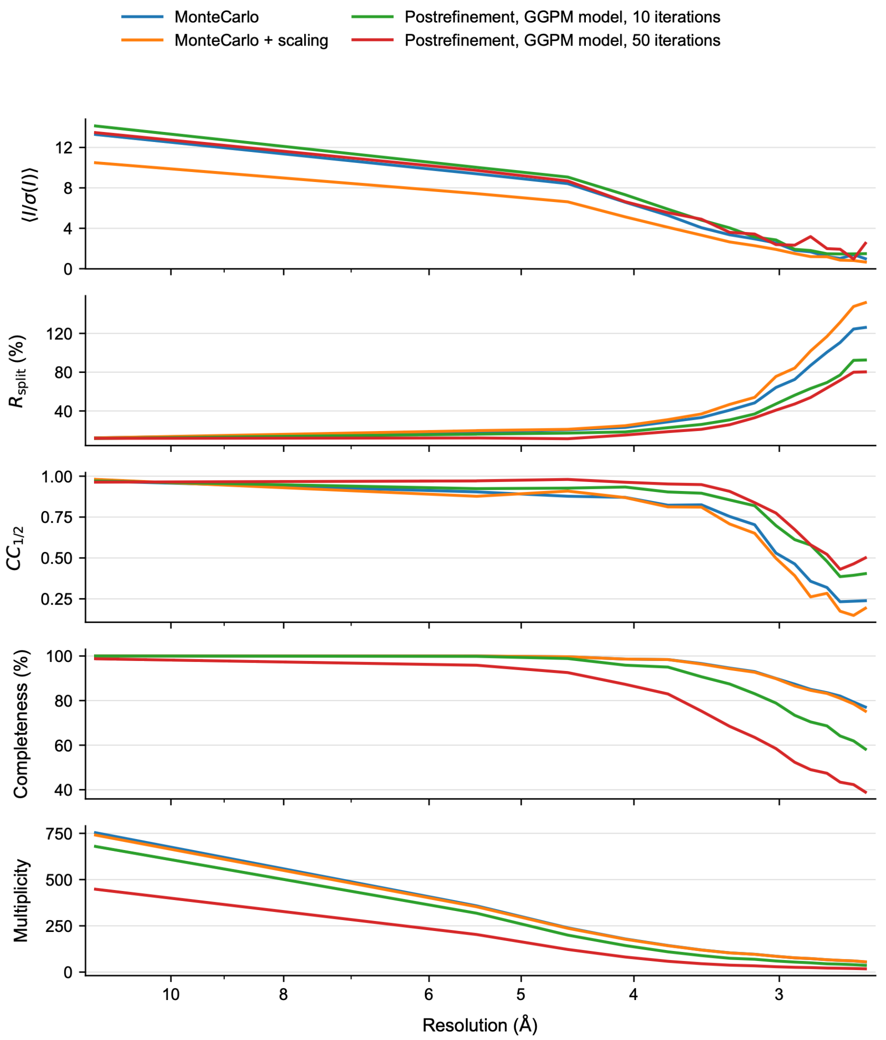


**Supplementary Fig. 3:** Comparison of merging strategies for SerialED data. The SerialED data were merged using several approaches to identify the strategy that best preserved data quality at high resolution. Monte Carlo averaging of observations without second-pass scaling provided the best overall merging strategy for the Cry11Aa SerialED data. Post-refinement using the GGPM model improved the quality indicators of the merged data, albeit at the expense of completeness and multiplicity.

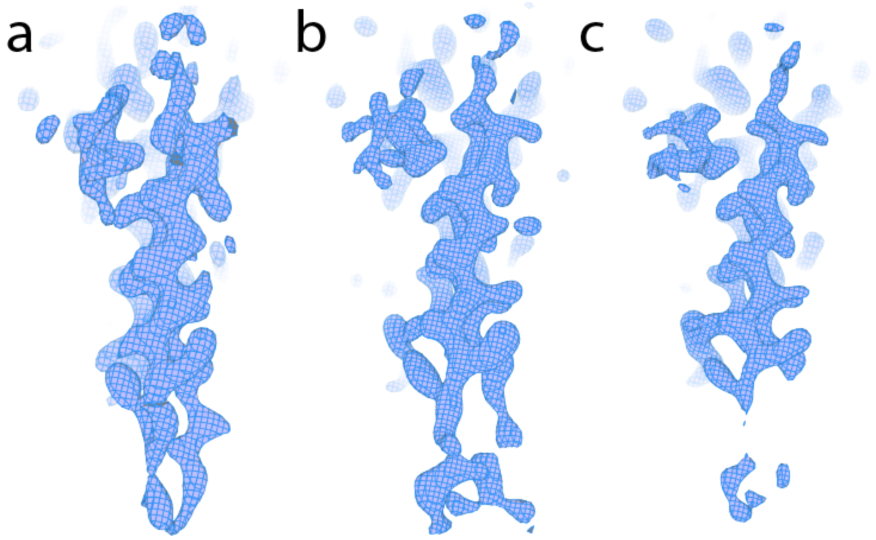


**Supplementary Fig. 4 Comparison of density maps around the loop and beta turn in the missing data direction.** a) MicroED. b) SerialED. c) SFX.

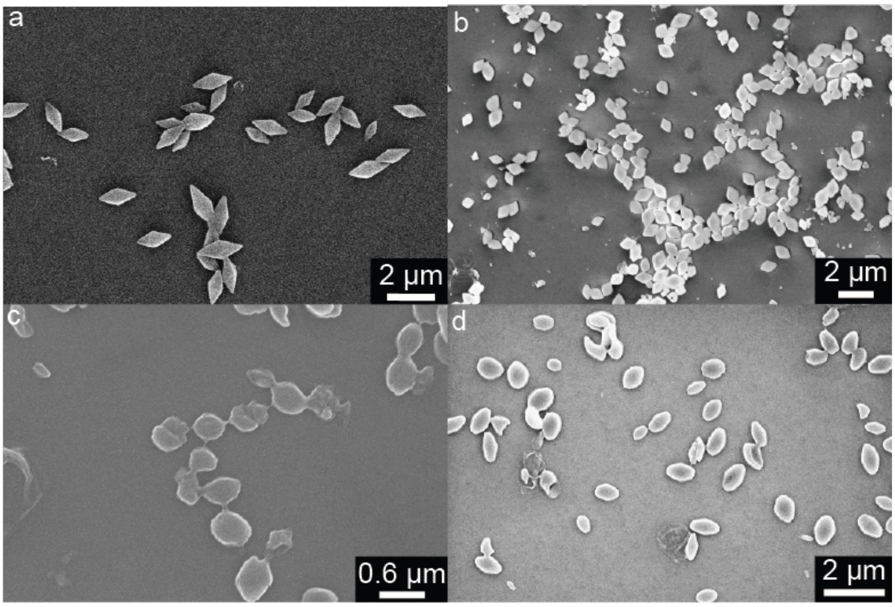


**Suplementary Fig. 5: Scanning electron micrographs of *Bt* grown nanocrystals. Crystals were deposited on glass coverslides** (a) *Bta* Cry1Ae; (b) *Bti* Cyt1A; (c) C11AB; (d) *Bti* Cry11Aa.

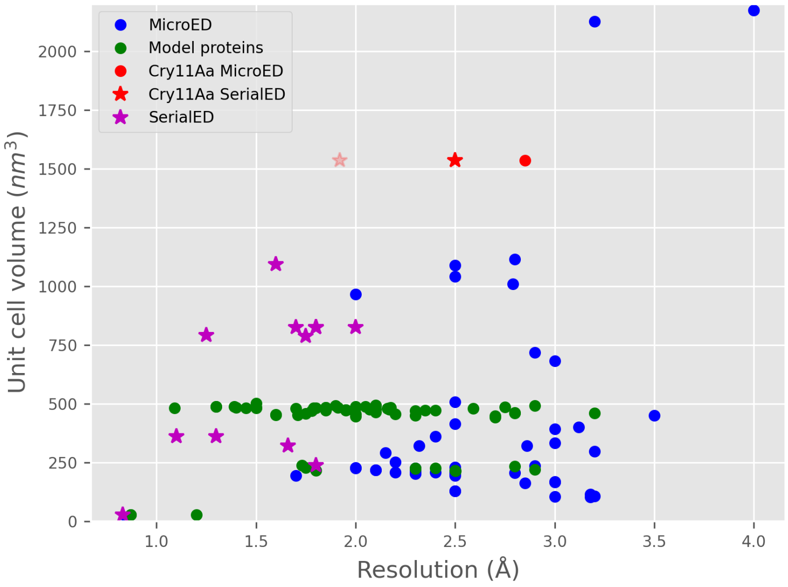


**Supplementary Fig. 6. Comparison between the unit cell volume and achieved resolution for all protein structures solved by MicroED and SerialED currently in the PDB.** The red circle and star indicates the Cry11Aa structure solved by MicroED and SerialED respectively in this work, amongst the largest unit cell size probed by either technique. The transparent red star shows the anisotropic diffraction limit of the Cry11Aa SerialED data showing the potential for further improvement. Magenta stars represent structures solved by SerialED showing the potential for improvement in resolution with this technique, particularly for crystals with larger unit cells.

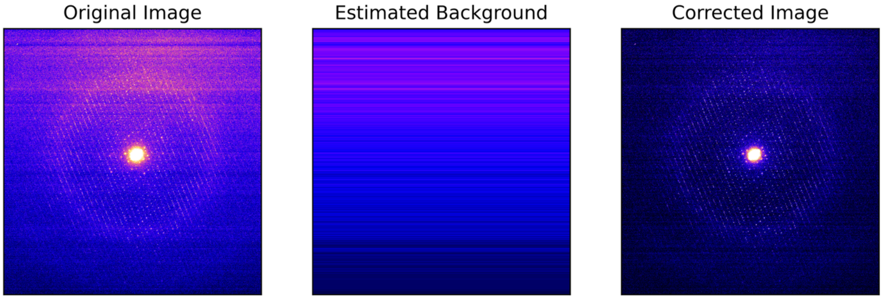


**Supplementary Fig. 7. Additional background correction for low fluence diffraction images captured on the K2**. The additional fluctuations in intensity are due to a line by line gain offset. This is estimated by calculating the median of each line in regions where there is now Bragg reflections and subsequently subtracted from the original image to produce a corrected image.

**Supplementary Table 1: Key data collection parameters that define the detection limit of diffraction in an MicroED experiment of Cry11Aa.**

| | |
|---|---|
| Beam diameter (Å) | 8000 |
| Illuminated area (Å$^2$) | 50,265,482 |
| Fluence (e$^-$/Å$^2$) | 4.5 |
| Total electrons passing through sample | 226,194,669 |
| Unit cell volume (Å$^3$) | 1,535,805 |
| Illumination volume (Å$^3$) – assuming 175 nm thick crystal | 87,964,593,500 |
| Total unit cells illuminated | 57,275 |

**Supplementary Table 2: Data collection statistics for MicroED data at lowered fluence**

| | Cry11Aa, MicroED<br>2 e$^-$/Å$^2$ total fluence |
|---|---|
| **Data collection** | |
| Wavelength (Å) | 0.025 |
| Space group | I222 |
| Cell dimensions | |
| *a*, *b*, *c* (Å) | 58.1, 155.6, 170.3 |
| α, β, γ (°) | 90, 90, 90 |
| Probe size (μm) | 0.8 |
| Number of crystals | 6 |
| Number of indexed patterns | 180 |
| Number of merged images | 180 |
| Resolution (Å) [a] | 85.41-3.96<br>(4.03-3.96) |
| Number of observations | 88171 (1200) |
| Number of unique reflections | 4710 (167) |
| I/σI [a] | 7.1 (1.9) |
| Rpim | 0.125 (0.364) |
| Rmerge | 0.392 (0.835) |
| $CC_{1/2}$ [a] | 0.942 (0.316) |
| Completeness (%)[a] | 66.50 (49.90) |
| Multiplicity [a] | 18.7 (7.2) |
